# Strategies for Overcoming the Purification Challenges of Meis1 and HoxA9 and Their Functional Characterization

**DOI:** 10.64898/2026.10.09.757953

**Authors:** Ibukunoluwa Abigail Olaosebikan, Kevin C. Courtney, Hacer Karatas Bristow

## Abstract

Transcription factors are critical regulators of homeostasis, and disruption of their function is a common underlying mechanism in cancer. The homeobox Myeloid Ecotropic Integration site-1 (Meis1) and its binding partner Homeobox A9 (HoxA9) are among the most highly dysregulated transcription factors in cancer, particularly in Acute Myeloid Leukemia (AML). Upregulation of HoxA9 is observed in more than 50% of AML cases and is a predictor of poor prognosis. Meis1 forms a cooperative DNA complex with HoxA9 and is required for HoxA9-mediated leukemogenesis. However, the mechanism by which HoxA9 and Meis1 interact and form a complex with DNA is unknown. To study the molecular mechanisms behind the HoxA9-Meis1-DNA complex and support future development of inhibitors, we describe optimized procedures for the recombinant expression and purification of full-length Meis1 and HoxA9. The constructs with an N-terminal His-tag showed degradation, but the addition of a small ubiquitin-like modifier (SUMO)-fusion tag improved solubility and stability for both proteins. Conditions were optimized for SUMO cleavage to obtain tag-free Meis1 and HoxA9 with high purity. We showed that both purified proteins bound to DNA with nanomolar affinities using a fluorescence polarization (FP) binding assay. The SUMO-fusion HoxA9 showed a distinct binding profile compared to the tag-free HoxA9 and formed a cooperative complex with Meis1. The tag-free HoxA9 appeared to oligomerize under the assay conditions. To the best of our knowledge, this is the first report to visualize the HoxA9-Meis1 cooperative interaction in a high-throughput assay. Our work establishes a foundation for future studies to decipher the protein-protein interaction of Meis1-HoxA9 and the development of inhibitors.

## INTRODUCTION

Acute Myeloid Leukemia (AML) is a bone marrow cancer that stems from the abnormal proliferation of myeloid stem cells.^1^ Although various approaches exist for targeting AML, including chemotherapy, stem cell transplantation, and targeted therapies, at least twelve of which have been FDA-approved within the last decade, the prognosis remains poor, and the relapse rates are high.^2–4^ Therefore, novel therapeutics that specifically target the molecular driving factors of AML are needed.

One of the critical driving factors in AML is the cooperation of transcription factor Homeobox A9 (HoxA9) with cofactors Myeloid Ecotropic Viral Integration Site Homolog 1 (Meis1) and Pre-B-cell Leukemia Homeobox (Pbx). Meis1, Pbx, and HoxA9 (MPH9) form binary and ternary complexes (i.e., HoxA9-Meis1, Meis1-Pbx, HoxA9-Pbx, HoxA9-Meis1-Pbx) on their cognate DNA sequences at the enhancer regions to activate or downregulate target genes.^5–7^ Genome-wide studies have identified different target downstream genes associated with HoxA9/Meis1 activity, including FLT3, MYB, SOX4, ERG and MYC, which participate in cell signaling, proliferation, and maintenance of the leukemic state.^7–9^

MPH9 is highly expressed in hematopoietic stem and progenitor cells to maintain self-renewal, then carefully downregulated upon differentiation to mature blood cells.^10,11^ The MPH9 complexes, particularly HoxA9-Meis1, are important in leukemogenesis, as their co-expression induces three-times rapid onset of leukemia in murine models compared to HoxA9 alone.^12,13^ The aberrant overexpression of HoxA9 is detected in over 50% of AML cases and is associated with poor prognosis, treatment failure, and poorer patient outcomes.^14-16^ The importance of HoxA9-Meis1 interactions extends across different AML subtypes. Deregulation of HoxA9 and Meis1 is particularly characteristic of leukemias containing KMT2A (MLL1) rearrangements (MLL_r_), NPM1c mutations, and HoxA9-NUP98 rearrangements, indicating that diverse genetic abnormalities can initiate AML due to the HoxA9/Meis1 transcriptional program.^17,18^

HoxA9 belongs to the HOX family of transcription factors that comprise 39 Mammalian homologs grouped into four clusters (A-D) and 1-13 paralogs.^19^ HoxA9 uses a homeodomain to bind DNA and a hexapeptide - also known as a pentapeptide or tryptophan motif - that contains a single conserved tryptophan residue to interact with Pbx (Figures 1A-1B). Meis1, one of the three Meis homologs, is composed of three domains: a homology region (HR-1 and HR-2) that forms protein-protein interactions with Pbx, a three-amino-acid extension loop (TALE) homeodomain that binds the DNA, and an unstructured transactivation domain at the C-terminus (Figures 1A-1B).^20,21^

**Figure 1.**
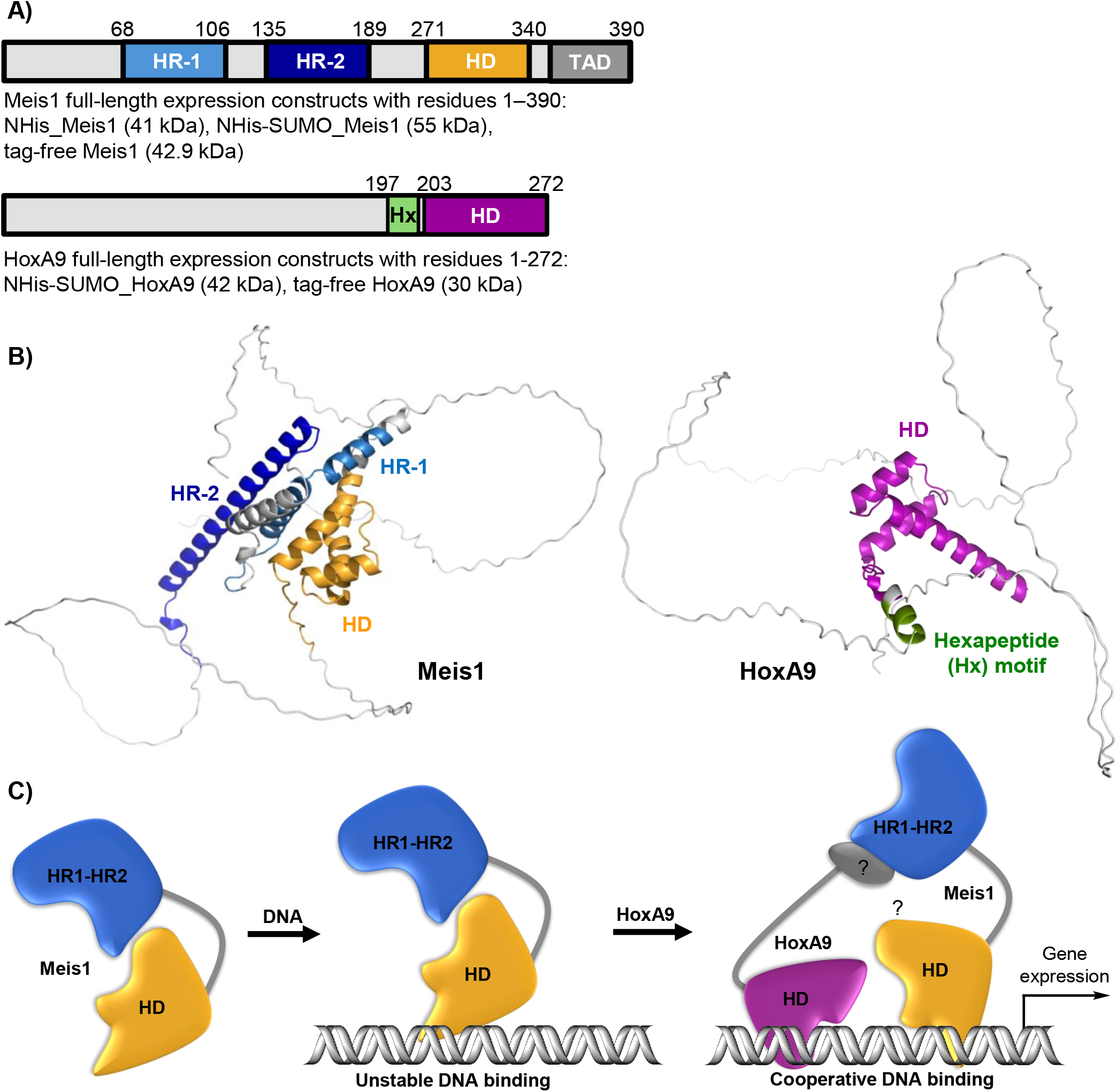
Structural organization of Meis1 and 390 HoxA9. A) Expression constructs used in the study. HR: homology region, HD: homeodomain, TAD: Transactivation domain, Hx: hexapeptide motif B) Meis1 full-length expression constructs with residues 1‒390: NHis_Meis1 (41 kDa), NHis-SUMO_Meis1 (55 kDa), tag-free Meis1 (42.9 kDa) Structures of Meis1 and HoxA9 as predicted with AlphaFold3. C) Proposed interaction of Meis1 with the DNA and HoxA9. Meis1 forms an unstable complex with the DNA, and an interaction with HoxA9 enhances Meis1 binding to the DNA. The mechanism of this cooperative interaction is unknown.

Meis proteins have been reported to form an unstable complex with DNA as detected in electrophoretic mobility shift assay (EMSA) experiments.^21,22^ This instability has been attributed to its HR domain repressing the homeodomain interaction with DNA, as the deletion of HR-2 in Meis2 reinstates transcription activation.^23^ HoxA9, which can interact with Meis1 in the presence or absence of DNA, significantly enhances Meis1 binding to DNA (Figure 1C).^21,24^ Furthermore, HoxA9 is a prerequisite for an association between Meis1 and chromatin as HoxA9 degradation leads to rapid loss of Meis1 from chromatin.^8^ Despite the importance of their cooperation in DNA binding and leukomogenesis,^25–27^ the details of the molecular interaction between Meis and HoxA9 are yet to be determined, partly due to limited in vitro studies examining their interactions. In general, purification of recombinant transcription factors can be challenging due to large intrinsically disordered regions, which often compromise protein stability and solubility. This technical challenge limits the biochemical characterization needed to define the molecular details of these interactions.

In this study, we have established, for the first time, optimized procedures for recombinant expression and purification of full-length Meis1 and HoxA9 tag-free proteins to investigate their individual and cooperative interaction with DNA. The N-terminal His-tag construct showed degradation during affinity purification, but the addition of a small ubiquitin-like modifier (SUMO)-fusion tag enhanced stability and solubility. We have optimized the conditions for SUMO cleavage to obtain Meis1 and HoxA9 with high purity and moderate yields. To the best of our knowledge, this is the first study showing that Meis1 and HoxA9 full-length proteins can be purified without a tag. We next showed that these purified proteins bind to the DNA in a fluorescence polarization (FP) binding assay. Lastly, we have shown that the cooperative interaction of Meis1 with HoxA9 can be visualized in the FP assay. This work provides a foundation for future studies to elucidate the Meis1-HoxA9 protein-protein interaction and develop inhibitors.

## RESULTS AND DISCUSSION

### The purification of NHis-Meis1 full-length leads to degradation

Meis1 is alternatively spliced into four isoforms, Meis1a‒d.^28^ We selected the Meis1a isoform (390 amino acids) for this study because its interaction with HoxA9 and role in HoxA9-mediated leukemogenesis is well established.^24,29,30^

We first expressed the NHis-Meis1 construct (Table S1) in *E. coli* via autoinduction.^31^ The cells were resuspended in Buffer-1 (pH 7.3, 25 mM Tris, 100 mM glycine, and 10% glycerol), lysed, purified over affinity chromatography, and the eluates were analyzed by sodium dodecyl sulfate-polyacrylamide gel electrophoresis (SDS-PAGE), showing a faint NHis-Meis1 band (44.2 kDa, purple arrow) along with numerous lower-molecular-weight, possibly degraded protein bands (Figure 2A). We then changed the buffer composition to Buffer-2 (pH 8.0, 50 mM Tris, 300 mM NaCl, 5% glycerol). However, degradation persisted, as shown in Figure 2B. Lowering the buffer pH to 7.5 (Buffer-3: pH 7.5, 50 mM Tris, 300 mM NaCl, 5% glycerol) also did not eliminate degradation (Figure 2C); hence, further optimization was needed.

**Figure 2.**
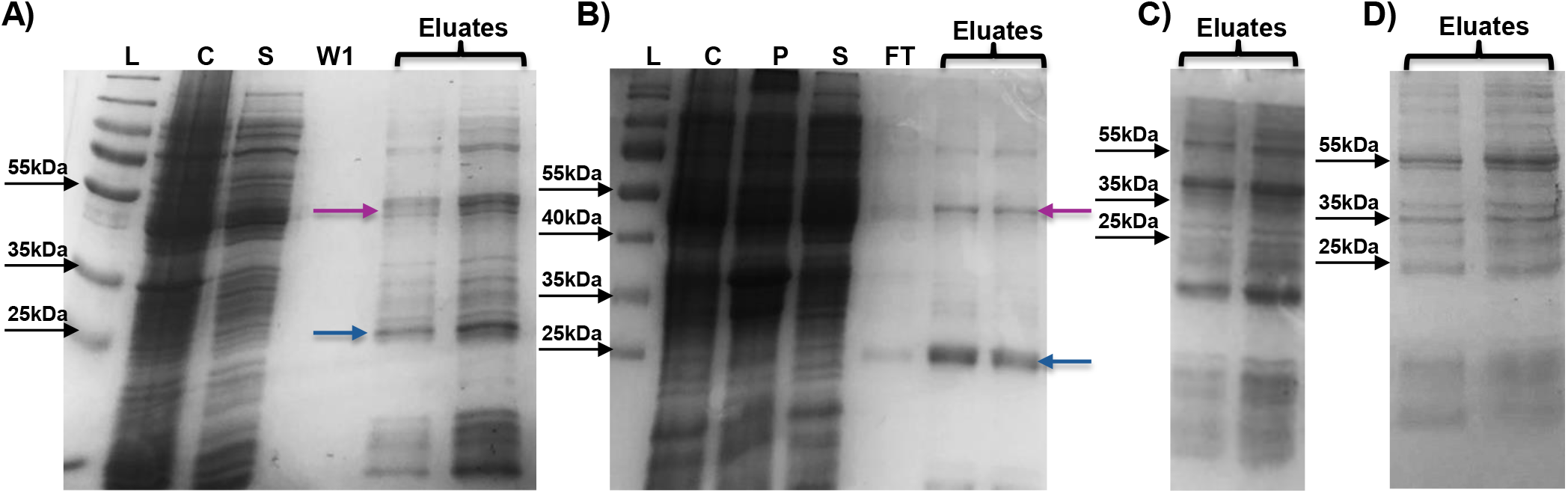
SDS-PAGE analysis of NHis-Meis1 purification. A) Degradation occurs during the purification of Meis1 (purple arrow), with a major degraded band of approximately 25 kDa (blue arrow). Changing B) the buffer composition entirely or C) pH from 8.0 to 7.5 did not reduce degradation. D) Reducing the incubation time with the beads from overnight to 4 hours reduced the intensity of the degraded bands. L-ladder, C-crude, S-supernatant, P-pellet, W1-wash.

Next, we tested whether Meis1 degraded during prolonged overnight incubation with the affinity beads, as used in Figures 2A‒2C, which could expose the protein to host proteases. After 4 hours of loading and under the same buffer conditions as in Figure 2B (i.e., Buffer-2), the degraded bands were still present but with reduced intensity compared to overnight loading (Figure 2D). While shortening the loading time reduced Meis1 degradation, it was still insufficient to obtain full-length Meis1. Since the attempts to optimize NHis-Meis1 purification conditions failed, we tested whether incorporating a stabilizing additive could help to improve the protein’s stability during purification. Accordingly, sucrose (2.5 mM), a stabilizing osmolyte, was added to Buffer-2 throughout the purification. We also reduced the loading duration to 2 hours. Although a faint full-length Meis1 band was observed in the elution fractions, degradation bands were still present, indicating that the addition of sucrose alone was insufficient to prevent protein degradation (Figure 3A). Further analysis of the wash fractions showed a second band, possibly a host protein, with a molecular weight close to that of Meis1 (Figure 3B).

**Figure 3.**
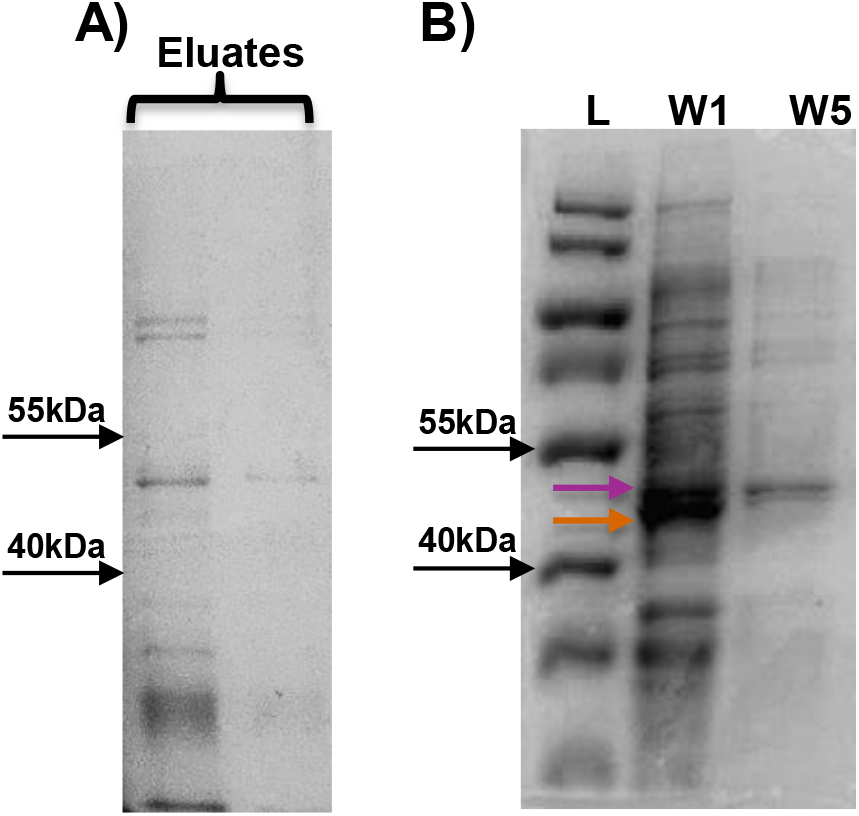
The addition of 2.5 mM sucrose as a stabilizing additive to the buffer does not reduce degradation. SDS-PAGE analysis reveals the Meis1 band in A) eluates and B) wash fraction (W1), with an additional band (orange arrow) showing a protein close to the size of Meis1 (purple arrow).

### IPTG induction improves the recombinant expression of Meis1

The additional band seen in the wash fractions (Figure 3B) suggested that the thick protein bands observed in crude, pellet, and supernatant upon autoinduction (Figures 2A-2B) might not solely belong to Meis1. Therefore, we performed IPTG-induced test expressions with a range of concentrations (0.1‒ 1.0 mM) and induction times (2, 4, and overnight) to determine Meis1 expression upon induction (Figure 4). The cells were centrifuged and lysed in the presence of SDS and analyzed with SDS-PAGE. A concentration of 0.4 mM IPTG yielded the strongest Meis1 band at all time points compared to all the other concentrations (Figure 4A), as determined by calculating the band intensities with ImageJ (Figure S1). Next, we performed a small-scale expression (i.e., 1.5 mL) with 0.4 mM IPTG induction for 4 hours, followed by test purification. SDS-PAGE analysis of the eluates showed a clear Meis1 band, indicating that the IPTG induction helped to recover the intact protein (Figure 4B). Although we observed some possible degradation bands, the intensity was lower than that observed for autoinduction under the same lysis and purification conditions (Figures 3A and 4B). Next, we tested a larger-scale Meis1 expression (1L culture volume) using the optimized induction conditions. The protein was purified by affinity (Figure 4C) followed by SEC, which revealed a broad peak suggesting a mixture of heterogeneous species. Further analysis of the combined SEC fractions confirmed the presence of Meis1; however, it was not pure (Fraction 2−4, Figure 4D). This result indicated a necessity for developing another Meis1 construct that would improve purity, stability, and solubility. Therefore, we designed a SUMO fusion construct.

**Figure 4.**
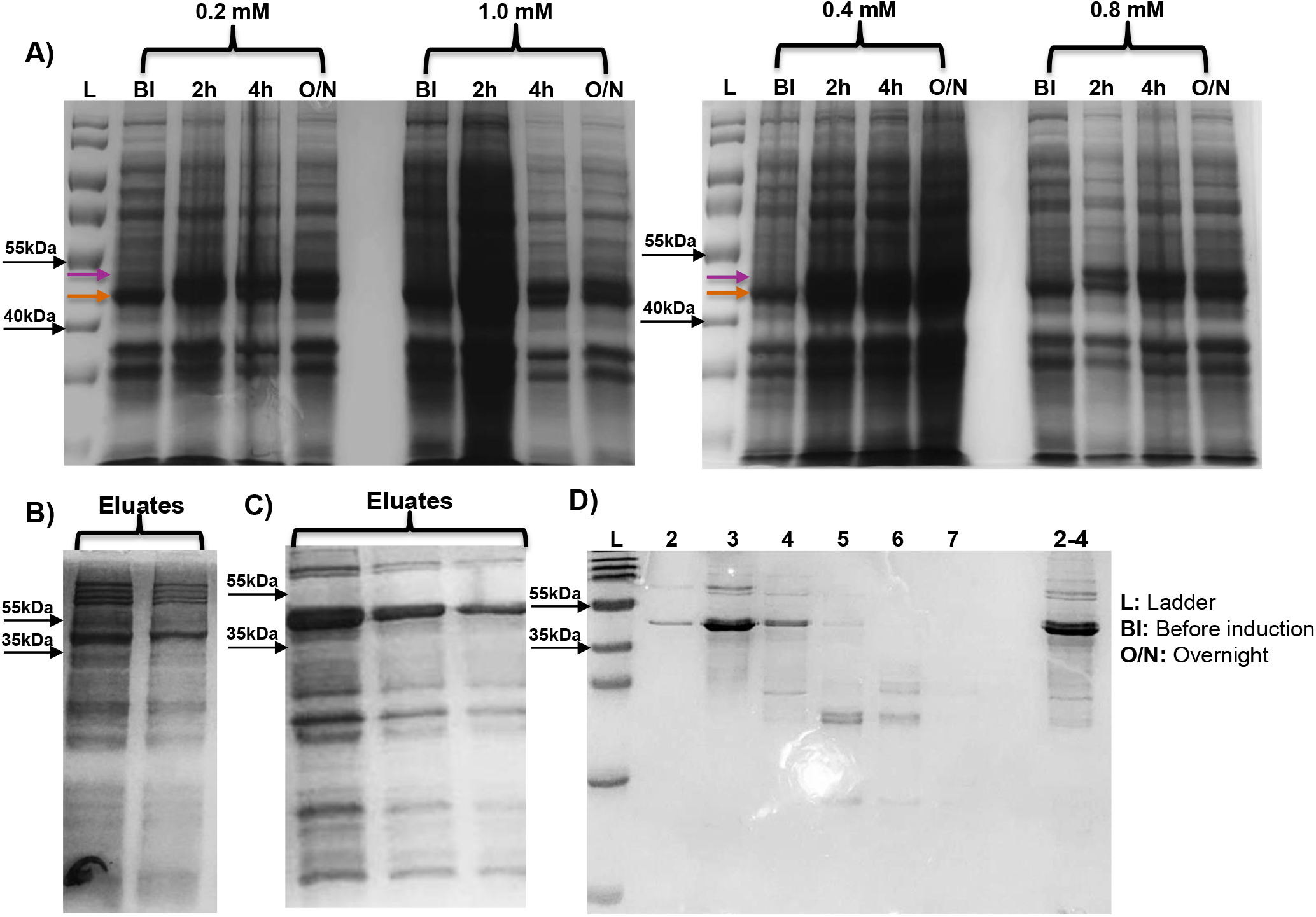
Optimization of Meis1 expression and purification. A) 0.2−1 mM IPTG-induced test expression at different time points before and after induction. The cells were induced at 37 °C for 2 and 4 hours and overnight at 30 °C. The host cell expresses a protein (orange arrow) with a similar size as Meis1 (purple arrow). B) 0.4 mM IPTG-induced test-expression-purification. C) 1L cell culture induced with 0.4 mM IPTG yields a thick Meis1 band and a low amount of degraded bands. D) Fractions (2‒7) collected during SEC and the concentrated samples of fractions 2‒4.

### SUMO-fusion stabilizes Meis1 and reduces degradation

SUMO fusions are commonly utilized to enhance protein solubility while preserving structural stability.^32^ Furthermore, the SUMO tag can be cleaved by a protease, such as *human* SENP2, which recognizes a Gly-Gly motif,^33^ to obtain a tag-free protein.^24,25^ Accordingly, we designed a new Meis1 construct with a cleavable His-SUMO tag at the N-terminus (Table S1).

We initially performed a small-scale (1.5 mL) expression using 0.4 mM and 0.8 mM IPTG induction for 2 h, 4 h, or overnight. The cell culture was centrifuged, lysed with SDS loading dye, and analyzed via SDS-PAGE (Figure 5A). A strong band at 55 kDa corresponding to His-SUMO_Meis1 was visible after 2 h induction with both IPTG concentrations, indicating successful expression. Induction with 0.4 mM IPTG for 2 hours yielded the strongest band (Figure 5A) among all other conditions and therefore was selected for further studies.

**Figure 5.**
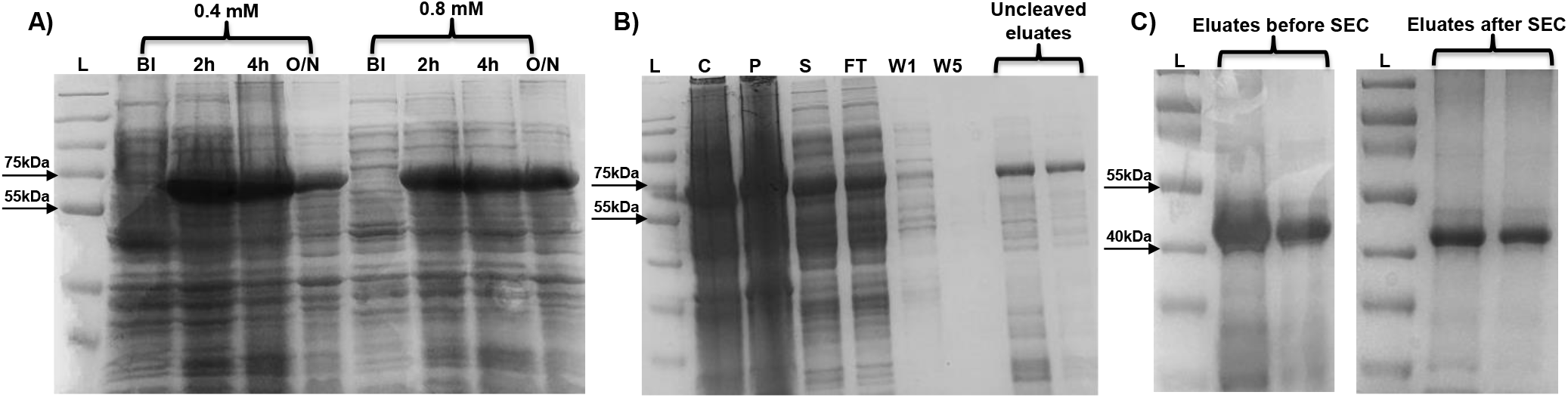
SUMO tag stabilizes Meis1 full-length. SDS-PAGE analysis of A) IPTG-induced cell test expression samples collected at different time points before induction and post-induction (2 and 4 hours after induction at 37 °C, as well as overnight at 30 °C). B) 0.4 mM IPTG-induced test-expressed and purified samples: C: crude, P: pellet, S: supernatant, W1 and W5: wash 1 and 5 respectively, F: flow-through, and eluates. C) Cleaved eluates obtained before and after SEC.

When a test affinity purification was performed on the test expression sample, the crude, pellet, supernatant, and eluates again showed the presence of the 55 kDa protein corresponding to NHis-SUMO_Meis1 (Figure 5B). We next scaled up the expression to a 1L culture volume with the optimized conditions (i.e., induction with 0.4 mM IPTG for 2 h at 37 °C). The cells were lysed and centrifuged, and the supernatant was loaded onto the beads overnight. Then, the beads were washed, resuspended in Buffer-2 with 2-mercaptoethanol (4 mM) and SUMO protease SENP2, and incubated overnight at 4 °C. The mixture was centrifuged to collect the cleaved Meis1 without the tag (42.9 kDa) in the supernatant, which was subsequently purified by SEC. The SDS-PAGE analysis of the eluates before and after SEC showed the presence of tag-free Meis1 (Figure 5C). In conclusion, the SUMO-fusion approach yielded tag-free Meis1 with high stability and purity.

### NHis-SUMO_HoxA9 full-length expression and purification

Initial attempts to purify HoxA9 full-length protein with a His-tag at the C-terminus failed with a similar degradation problem as observed with Meis1 (results not shown). Having established a workflow for Meis1 with a cleavable SUMO tag, we applied the same methodology to express and purify HoxA9 full-length (residues 1-272). Induction with 0.4 mM IPTG for 4 hours produced the strongest band corresponding to His-SUMO_HoxA9 (42 kDa) compared to the other conditions (Figure 6A, green arrow). On the other hand, in the next step, on-resin SENP2 treatment did not efficiently release HoxA9 into the solution (lanes C1-C2, Figure 6B). To assess whether HoxA9 is retained on the resin, we eluted the remaining bound proteins with imidazole (lane E-imi, Figure 6B), which revealed bands for HoxA9 (30 kDa, blue arrow) and His-SUMO (12 kDa, pink arrow). The latter is expected to be bound to the cobalt resin because of the His-tag. Retention of HoxA9 is possibly due to an internal poly-histidine sequence located at the N-terminus of HoxA9 (residues 84-89).

**Figure 6.**
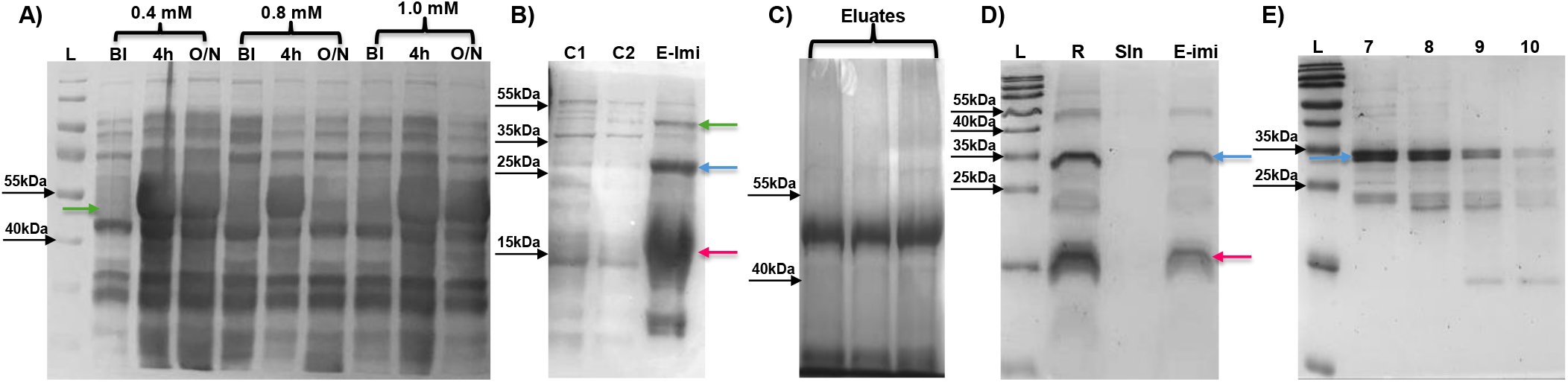
SDS-PAGE analysis of NHis-SUMO_HoxA9 full-length. A) IPTG-induced test expression samples collected before induction and post-induction (induction for 2 and 4 hours at 37 °C, overnight at 30 °C). B) Cleavage samples from SENP2 treatment. C1-C2: Supernatant from SENP2 treatment. E-imi: imidazole-eluted samples. The His-SUMO_HoxA9 (green arrow, 42.2 kDa), HoxA9 (blue arrow, 30 kDa), and His-SUMO (pink arrow, 12 kDa) are indicated. C) Eluates of NHis-SUMO_HoxA9 full-length (42 kDa). D) Cleaved products of NHis-SUMO_HoxA9 after SENP2 treatment. R: SENP2-treated reaction mixture, Sln: solution after loading, E-Imi: imidazole-eluted fraction. E) SEC fractions of the tag-free HoxA9.

Because the on-resin cleavage/purification method for HoxA9 failed, we decided to elute the protein as a fusion, purify it by SEC or dialysis, and then cleave the SUMO tag off-resin. However, our initial attempt to SEC-purify His-SUMO_HoxA9 using Buffer-2 failed due to protein aggregation (results not shown). Therefore, we changed the buffer pH to 6.5 and added arginine as an additive to prevent aggregation, leading to successful affinity and SEC purification, as shown in Figure 6C. We used the same buffer composition for the rest of the optimization procedure.

Next, we treated the SEC-purified NHis-SUMO-HoxA9 with 0.02 equiv. of SENP2 protease overnight, followed by loading the reaction mixture onto the beads to trap the NHis-SUMO and obtain HoxA9 in solution. While two bands corresponding to HoxA9 (30 kDa) and His-SUMO (12 kDa) were detected in the reaction mixture (Figure 6D, lane R), HoxA9 was not in solution after the loading step (Figure 6D, lane Sln). Upon treating the beads with imidazole, we found that both NHis-SUMO and HoxA9 were bound to the beads (Figure 6D, lane E-Imi, pink and blue arrows, respectively). It is evident that SENP2 efficiently cleaved the SUMO tag; however, HoxA9 has again been retained on the resin. Therefore, we modified the procedure by dialyzing the fusion protein after affinity purification, cleaving the SUMO tag in solution, and purifying the reaction mixture with SEC. Fractions 7-8 showed the presence of a strong band near 30 kDa corresponding to tag-free HoxA9 (Figure 6E, blue arrow).

### Binding of Meis1 and HoxA9 to DNA

We next optimized an FP binding assay to determine Meis1 and HoxA9 binding to a fluorescently labeled 30-base pair double-stranded oligonucleotide containing Meis1 (<u>TGACAG</u>) and HoxA9 (TTTAC) binding motifs. Saturation binding curves showed that Meis1 binds to the DNA with *K*_d_ = 15.2 nM at 0 min, and the affinity increases slightly at 60 min (*K*_d_ = 10.6 nM) (Figure 7A). However, we observed a decrease in the signal span over time. Because the dynamic range in an FP assay correlates with the size and/or shape of the fluorophore-macromolecule complex^34^, this decrease in signal span suggests that Meis1 binding to DNA may interfere with its structural rigidity.

**Figure 7.**
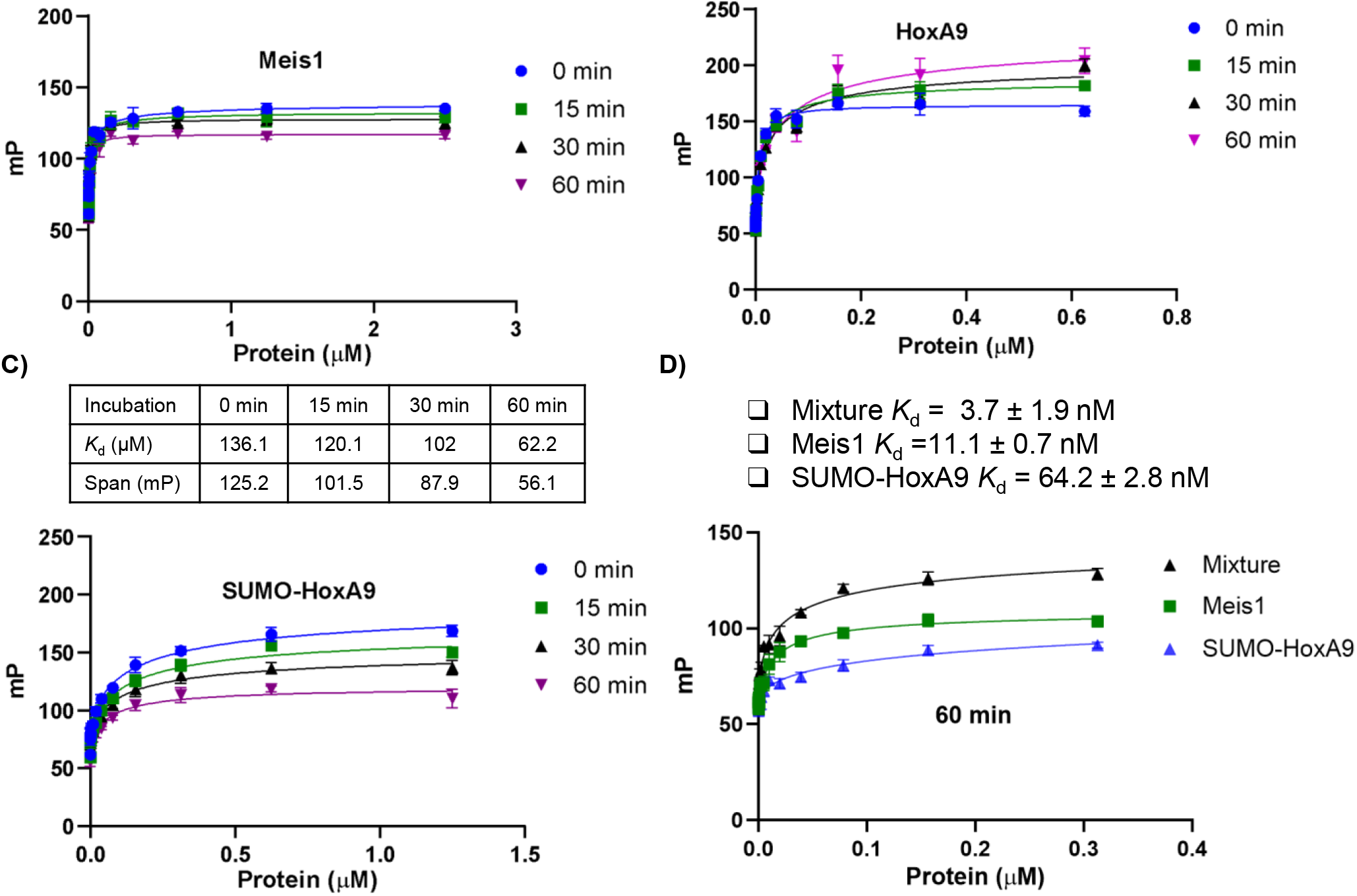
FP binding experiments. Saturation binding curves of A) Meis1, B) HoxA9, C) NHis-SUMO_HoxA9 over 60 min duration with a 5/6-FAM-labeled DNA oligonucleotide, i.e., 5-ccagatc<u>TGACAG</u>TTTTACGACagatctcc-3, carrying cognate binding sequences of <u>Meis1</u> and HoxA9 as underlined. D) A mixture of Meis1 and HoxA9 enhances the affinity and the signal span compared to individual proteins alone, indicating cooperative DNA binding. Imaging data at 60 min is shown.

As noted earlier, Meis has been shown to form an unstable complex with DNA, possibly because the HR 1-2 region restricts its binding to DNA (Figure 1C).^21^ Our study with AlphaFold3 modeling predicted a potential intramolecular interaction between HR1-2 and the homeodomain (Figure 1B), suggesting a possible structural basis for the unstable DNA-binding behavior of Meis1 reported in the literature.^21^ However, experimental structural studies are required to validate this predicted interaction.^18^

FP saturation binding experiments with tag-free HoxA9 (Figure 7B) showed a time-dependent increase in signal response and a 6-7-fold decrease in binding affinity (*K*_d_ = 7 nM at 0 min, *K*_d_ = 44 nM at 60 min). In general, the progressive increase in signal over time is reminiscent of oligomerization, which suggests that HoxA9 might be dimerizing under the assay conditions. Selected Hox proteins have been shown to homo- or heterodimerize with other Hox proteins; however, the physiological role of such complexes has not been deciphered.^35^

We next tested the fusion protein, SUMO-HoxA9, for binding to the FAM-labeled DNA and whether it would exert a similar binding profile as the tag-free HoxA9. The fusion protein binds to the DNA with *K*_d_ = 136 nM at 0 min, and the affinity increased to 62 nM at 60 min, while the DNA-protein complex signal decreased 2-fold (Figure 7C). We hypothesize that the NHis-SUMO_HoxA9 binding to the DNA might lead to a significant conformational change that allows the homeodomain to efficiently interact with the DNA. Unlike tag-free HoxA9, the SUMO-HoxA9 did not appear to be dimerized under the assay conditions and therefore was chosen for cooperative complex experiments.

Former studies with HoxA9 and Meis1 relied on EMSA, which is not high-throughput and requires the handling of delicate polyacrylamide gels.^6,21^ The FP assay, on the other hand, is a homogeneous assay and suitable for screening. Therefore, we tested whether our FP assay could detect a cooperative DNA binding with HoxA9 and Meis1. The Meis1 and SUMO-HoxA9 mixture (*K*_d_ = 3.7 nM) bound to the DNA with 3- and 20-fold higher affinity, respectively, compared to individual Meis1 (*K*_d_ = 11.1 nM) and SUMO-HoxA9 (*K*_d_ = 64.2 nM) proteins at 60 mins (Figure 7D). Additionally, the wider dynamic range of the mixture compared to either protein alone indicated a larger DNA complex, further supporting the formation of a HoxA9-Meis1 cooperative complex. To the best of our knowledge, this is the first report that a HoxA9-Meis1 cooperative interaction can be visualized in a throughput assay. On the other hand, the tag-free HoxA9 exerted no cooperation with Meis1 (Figure S3). A similar signal span and *K*_d_ values in the presence and absence of Meis1 suggest that an overlapping region in HoxA9 might be playing a role in interaction with Meis1 and homodimerization.

## Conclusions

Here we have shown a detailed optimization of recombinant expression and purification of transcription factors Meis1 and HoxA9. Both proteins are prone to degradation and are unstable when expressed with a His-tag at the N-terminus. Adding a soluble SUMO tag, however, improved the stability of both proteins. An internal poly-histidine sequence in HoxA9 prevented on-resin SUMO cleavage as the protein bound to the resin, which was resolved by off-resin SUMO cleavage followed by SEC purification. Next, we established an FP assay for Meis1 and HoxA9, which showed nanomolar affinities when bound to a DNA oligonucleotide that contains cognate DNA sequences. Meis1 forms a time-dependent complex with DNA, and this observation is consistent with the previous findings that Meis1/DNA complexes dissociate more rapidly than Meis1/HoxA9/DNA complexes. The SUMO-fusion HoxA9 showed a different binding profile compared to tag-free HoxA9 and formed a cooperative complex with Meis1 in the FP assay. This is the first report that the HoxA9-Meis1 cooperative interaction can be visualized in a high-throughput assay. Our work is a foundation for future studies to decipher the protein-protein interaction of Meis1-HoxA9 and the development of inhibitors.

## METHODS

### Protein constructs

Plasmids encoding full-length human Meis1 (1-390) and HoxA9 (1-272) as N-His-tagged and N-His-SUMO-tagged constructs were synthesized and subcloned by GenScript (Piscataway, NJ) into expression vectors: pET-28b (+) and pET-28a (+) under the control of the T7 promoter, respectively. The lyophilized plasmids were resuspended in deionized water to a final concentration of 40 ng/µl and chemically transformed into *E. coli* strain BL21 (DE3) competent cells.

### Protein expression and purification

Protein expression was achieved using either autoinduction or IPTG induction in lysogeny broth (LB) medium containing Kanamycin (50 µg/mL).

#### Autoinduction

A pre-culture was prepared by adding bacteria from a glycerol stock to LB medium with glucose (1%) and MgSO_4_ (24 mg/mL), then shaking overnight at 37 °C. The preculture was then transferred (1/400 ratio) to the culture medium with lactose (10 mg/mL) and incubated for 4 hours at 37 °C, then at 25 °C for 22 hours.

#### IPTG induction

The cells were grown at 37 °C to mid-log phase (0.6-0.8), and expression was induced with 0.1-1.0 mM IPTG (Sigma-Aldrich) for 4 hours at 37 °C. During the test expression (5 mL culture volume) the temperature was lowered to 30 °C and further shaken overnight.

#### Harvesting and purification

The cells were harvested by centrifugation at 4000 *g* for 25 mins at 4 °C; the supernatant was discarded, and the pellet was resuspended in a 35 mL lysis buffer (or 1.5 mL for test expression). EDTA-free protease inhibitor cocktail (Pierce, Thermo Scientific) and 2-mercaptoethanol (4 mM) were added to the resuspended cells which were sonicated for 5 min at 50% amplitude, 5 sec on/off with a 0.25” probe. The test expression sample in 1.5 mL lysis buffer was sonicated for 10 pulses (0.078” probe). The lysate was centrifuged at 15,000 *g* for 60 mins followed by affinity purification. The supernatant was loaded onto cobalt beads (HisPur Cobalt Resin, Thermo Scientific) overnight. Then the beads were washed 5-10 times with wash buffer (50 mM Tris-HCl, 300 mM NaCl, 5% glycerol, 5 mM imidazole, pH 8.0 or 6.5) and treated with elution buffer (50 mM Tris-HCl, 300 mM NaCl, 5% glycerol, 4 mM 2-mercaptoethanol, pH 8.0 or 6.5) at 4 °C containing SENP2 (overnight) or imidazole (30 mins). The eluates were further purified by SEC (ACTA Gold, Cytiva Superdex 75 Increase column of 10/300) equilibrated in 50 mM Tris-HCl, 300 mM NaCl, 5% glycerol, pH 8.0 or 6.5, at a flow rate of 0.8 mL/min. Peak fractions were collected, concentrated, and the concentration was quantified using a NanoDrop spectrophotometer and the Bradford assay. Protein samples were aliquoted, flash-frozen, and stored at -80 °C.

### DNA oligonucleotide

Double-stranded DNA oligonucleotide labeled with 5-carboxyfluorescein (5/6-FAM) was purchased from Integrated DNA Technologies (IDT). A 100 µM stock solution was prepared in 1x DNA resuspension buffer containing Tris-HCl 10 mM, NaCl 150 mM, EDTA 1 mM, DTT 0.1 mM, pH 7.5. The oligonucleotide was aliquoted and stored at -20 C.

**Table 1.**
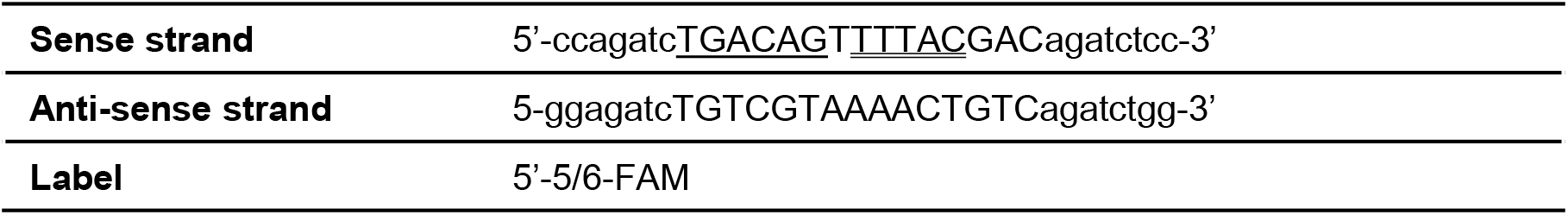
Sequence of the oligonucleotide used in the study. Cognate binding sequences for <u>Meis1</u> and HoxA9 are underlined.

### Saturation binding experiments

FP binding experiments were performed using a non-binding round-bottomed 384-well plate (Corning 4514). The assay buffer was 1/10 diluted from 10X buffer (10 mM HEPES, 100 mM NaCl, 5% glycerol, 2 mM EDTA, pH 7.5). To 5 µL of serially diluted protein, 5 µL of 10 nM FAM-labelled DNA solution was added. The plate was shaken at room temperature, and the FP was measured (Ex/Em: 485/530) using a microplate reader (TECAN Spark). The K_D_ was calculated using GraphPad Prism 10.

## Author Contributions

Ibukunoluwa Abigail Olaosebikan: Conceptualization, methodology, validation, formal analysis, investigation, writing - original draft; Kevin C. Courtney: Review and editing, supervision. Hacer Karatas Bristow: Methodology, supervision, writing - review & editing, project administration. All authors have read, commented on, and agreed to the submitted version of the manuscript.

## Data availability

All data generated and analyzed are available in the main text and the supplementary information (SI). For additional materials on construct sequences, protein purification SEC chromatogram and SDS-PAGE, as well as more fluorescence polarization data, please see the SI.

## Acknowledgement

This research was supported by start-up funds awarded to Hacer Karatas Bristow from the Department of Chemistry at West Virginia University. The authors are grateful to the Biochemistry Viral Core Facility at the Department of Biochemistry and Molecular Medicine in the Health Sciences Center at West Virginia University for providing access to the FPLC system used in this study.

