## Supporting information for "Strategies for Overcoming the Purification Challenges of Meis1 and HoxA9 and Their Functional Characterization"

| Construct | Residues | Sequence |
|---|---|---|
| **NHis_Meis1** | **1-390** | MHHHHHHAQRYDDLPHYGGMDGVGIPSTMYGDPHAARSMQPVHHLNHGPPLHSHQYPHT<br>AHTNAMAPSMGSSVNDALKRDKDAIYGHPLFPLLALIFEKCELATCTPREPGVAGGDVCSSES<br>FNEDIAVFAKQIRAEKPLFSSNPELDNLMIQAIQVLRFHLLELEKVHELCDNFCHRYISCLKGKM<br>PIDLVIDDREGGSKSDSEDITRSANLTDQPSWNRDHDDTASTRSGGTPGPSSGGHTSHSGD<br>NSSEQGDGLDNSVASPSTGDDDDPDKDKKRHKKRGIFPKVATNIMRAWLFQHLTHPYPSEE<br>QKKQLAQDTGLTILQVNNWFINARRRIVQPMIDQSNRAVSQGTPYNPDGQPMGGFVMDGQQ<br>HMGIRAPGPMSGMGMNMGMEGQWHYM |
| **NHis-SUMO_Meis1** | **1-390** | MHHHHHHMSDSEVNQEAKPEVKPEVKPETHINLKVSDGSSEIFFKIKKTTPLRRLMEAFAKRQ<br>GKEMDSLRFLYDGIRIQADQTPEDLDMEDNDIIEAHREQIGGAQRYDDLPHYGGMDGVGIPST<br>MYGDPHAARSMQPVHHLNHGPPLHSHQYPHTAHTNAMAPSMGSSVNDALKRDKDAIYGHP<br>LFPLLALIFEKCELATCTPREPGVAGGDVCSSESFNEDIAVFAKQIRAEKPLFSSNPELDNLMIQ<br>AIQVLRFHLLELEKVHELCDNFCHRYISCLKGKMPIDLVIDDREGGSKSDSEDITRSANLTDQP<br>SWNRDHDDTASTRSGGTPGPSSGGHTSHSGDNSSEQGDGLDNSVASPSTGDDDDPDKDK<br>KRHKKRGIFPKVATNIMRAWLFQHLTHPYPSEEQKKQLAQDTGLTILQVNNWFINARRRIVQP<br>MIDQSNRAVSQGTPYNPDGQPMGGFVMDGQQHMGIRAPGPMSGMGMNMGMEGQWHYM |
| **NHis_HoxA9** | **1-272** | MHHHHHHATTGALGNYYVDSFLLGADAADELSVGRYAPGTLGQPPRQAATLAEHPDFSPCS<br>FQSKATVFGASWNPVHAAGANAVPAAVYHHHHHHPYVHPQAPVAAAAPDGRYMRSWLEPT<br>PGALSFAGLPSSRPYGIKPEPLSARRGDCPTLDTHTLSLTDYACGSPPVDREKQPSEGAFSE<br>NNAENESGGDKPPIDPNNPAANWLHARSTRKKRCPYTKHQTLELEKEFLFNMYLTRDRRYEV<br>ARLLNLTERQVKIWFQNRRMKMKKINKDRAKDE |
| **NHis_SUMO_HoxA9** | **1-272** | MHHHHHHMSDSEVNQEAKPEVKPEVKPETHINLKVSDGSSEIFFKIKKTTPLRRLMEAFAKRQ<br>GKEMDSLRFLYDGIRIQADQTPEDLDMEDNDIIEAHREQIGGATTGALGNYYVDSFLLGADAA<br>DELSVGRYAPGTLGQPPRQAATLAEHPDFSPCSFQSKATVFGASWNPVHAAGANAVPAAVY<br>HHHHHHPYVHPQAPVAAAAPDGRYMRSWLEPTPGALSFAGLPSSRPYGIKPEPLSARRGDC<br>PTLDTHTLSLTDYACGSPPVDREKQPSEGAFSENNAENESGGDKPPIDPNNPAANWLHARST<br>RKKRCPYTKHQTLELEKEFLFNMYLTRDRRYEVARLLNLTERQVKIWFQNRRMKMKKINKDR<br>AKDE |

***Table S1. The sequences of the protein constructs used in the study.***

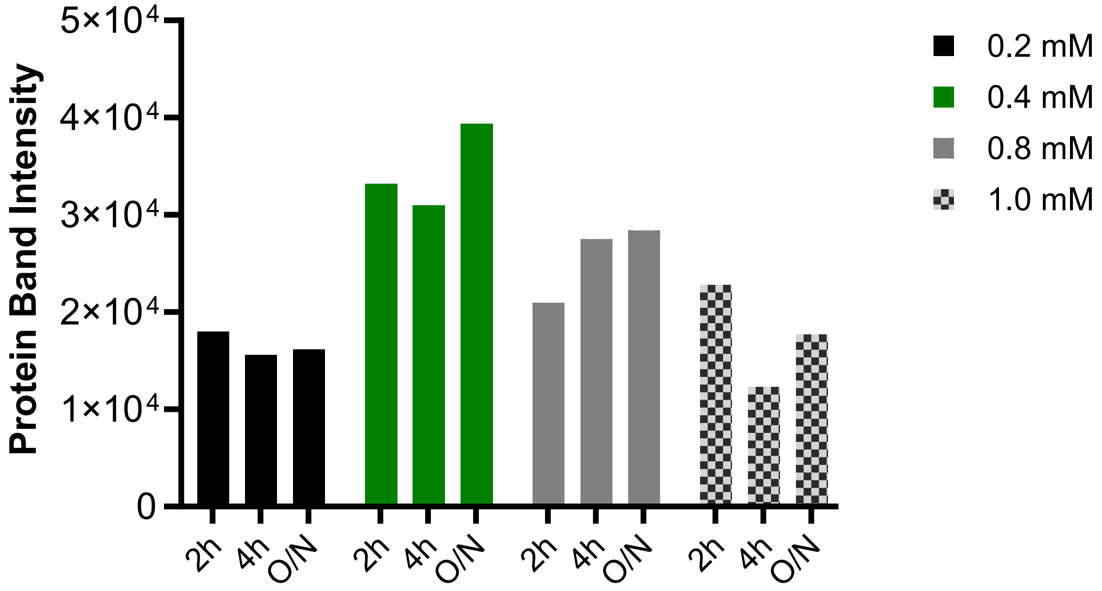


***Figure S1. Quantification of IPTG-induced NHis-Meis1 protein bands.*** *The intensity of the protein bands on the SDS-PAGE gel was measured using ImageJ software.*

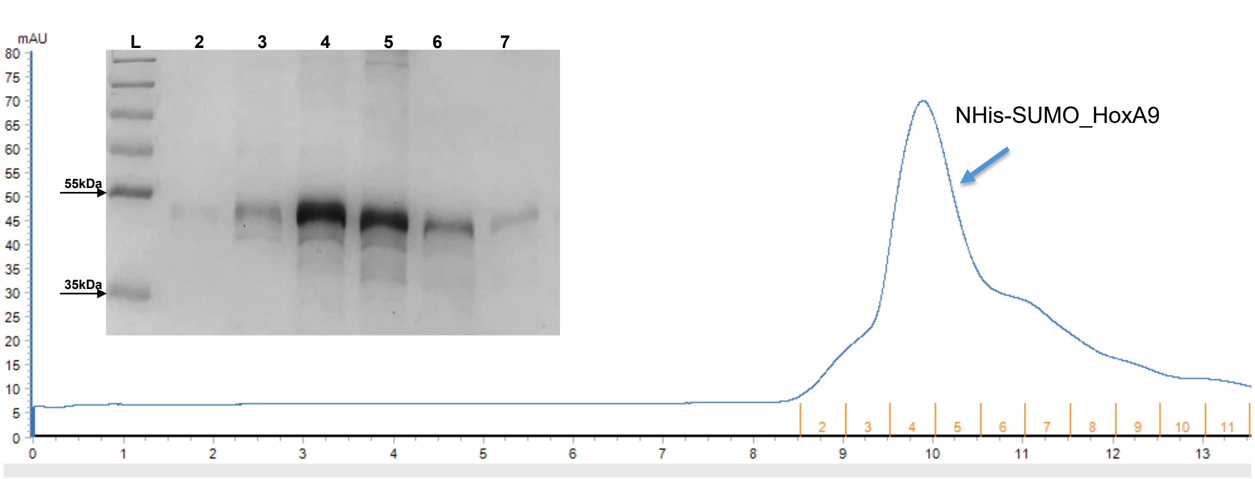


***Figure S2. SEC purification of NHis_SUMO-HoxA9.*** *SEC chromatogram revealed a narrow peak, which was analyzed by SDS-PAGE. Fractions 4 and 5 correspond to NHis-SUMO_HoxA9 (42 kDa).*

| Incubation | Meis1 | HoxA9 | Mixture |
|---|---|---|---|
| $K_d$ (nM) | 18 | 47 | 42 |
| Span (mP) | 54 | 175 | 172 |

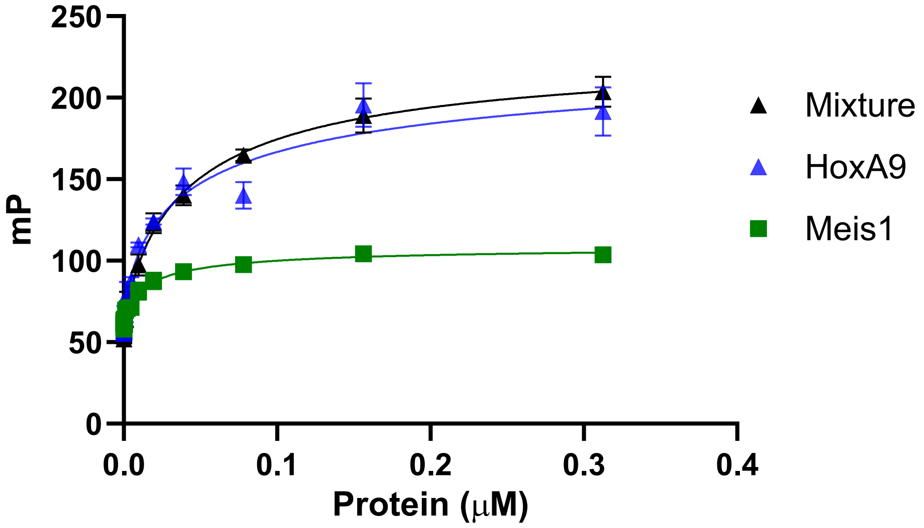


***Figure S3. FP binding experiments with tag-free proteins***. *Cooperative DNA binding of Meis1 and HoxA9 with a 5/6-FAM-labeled DNA oligonucleotide, i.e., 5-ccagatcTGACAGTTTTACGACagatctcc-3, carrying the cognate binding sequences of Meis1 and HoxA9, as underlined.*
